# Tau-mediated synaptic dysfunction is coupled with HCN channelopathy

**DOI:** 10.1101/2020.11.08.369488

**Authors:** Despoina Goniotaki, Francesco Tamagnini, Luca Biasetti, Svenja-Lotta Rumpf, Claire Troakes, Saskia J. Pollack, Shalom Ukwesa, Haoyue Sun, Louise C. Serpell, Wendy Noble, Kevin Staras, Diane P. Hanger

**Affiliations:** Department of Basic and Clinical Neuroscience, Institute of Psychiatry, Psychology & Neuroscience, King’s College London, UK; School of Pharmacy, University of Reading, Reading, UK; Sussex Neuroscience, School of Life Sciences, University of Sussex, UK

## Abstract

Progressive neurodegeneration in tauopathies is mediated through an elusive mechanism. Here, we show that hyperpolarization-activated cyclic nucleotide-gated (HCN) channels are functionally linked to disease-associated abnormalities in tau. Selective rises in the proportion of HCN-positive neurons are detected both in post-mortem human brain from Alzheimer’s disease and in the Tau35 mouse model of tauopathy. Tau35 mice develop progressive abnormalities including increased phosphorylated tau, enhanced HCN channel expression and decreased dendritic branching, as well as reduced synapse density that is accompanied by vesicle clustering defects. Notably, altered spine density and increased HCN channel expression in Tau35 neurons correlates with functional abnormalities in network properties, including enhanced hyperpolarization-induced membrane voltage ‘sag’ and changes in the frequency and kinetics of spontaneous excitatory postsynaptic currents. Our findings are consistent with pathological changes in tauopathies impacting on HCN channels to drive network-wide structural and functional synaptic deficits, providing new targets for therapeutic intervention.

## Introduction

Synapses are the principal structural and functional units that enable information signaling between neurons. Although synaptic dysfunction is an early correlate of dementia, the pathophysiology of aberrant synaptic signaling remains poorly understood (*1*). In human tauopathies and in mouse models of disease, accumulation of phosphorylated and truncated tau, altered tau processing and aberrant tau localization occur in parallel with reductions in presynaptic protein expression, synapse density and synaptic function, suggesting a causal role for tau in disease pathogenesis (*2–4*).

Voltage-gated hyperpolarization-activated cyclic nucleotide-gated (HCN) channels 1-4 are a family of non-selective cation channels that are emerging as central regulators in neuronal signaling (*5*). HCN channels are involved in multiple synaptic processes including the regulation of membrane resistance, the setting of intrinsic membrane excitability, the generation of synaptic potentials, and synaptic vesicle exocytosis (*6–8*). Notably, HCN channels selectively associate with regulatory proteins in dendritic spines to drive fine-tuning of network connectivity (*9*). HCN channel activation generates an inward-rectifying (I_h_) current, which reduces dendritic summation and synaptic vesicle release through interaction with calcium channels (*10–12*). For example, in adult entorhinal cortex, increased HCN channel expression dampens spontaneous synaptic vesicle release (*12*). In parallel, mis-regulation of HCN channels can result in either gain-of-function or loss-of-function in neurological disorders (*13*) and HCN1 is implicated in disease progression in transgenic animal models of Alzheimer’s disease (AD) (*14–16*). These wide-ranging roles of HCN channels in regulating key structure-function properties in neurons position them as potentially important substrates to explain the development and progression of tauopathies.

Here we investigate the link between tau abnormalities and HCN channels. We demonstrate an imbalance of HCN channel expression both in post-mortem AD human brain and in Tau35 mice, a progressive tauopathy model that exhibits increased tau phosphorylation and deposition of abnormal tau species in the brain (*17*). Strikingly, selective increases in HCN channels occur as Tau35 mice age, a change that is recapitulated both in dissociated cultured neurons as time *in vitro* increases, and in AD brain. In Tau35 mice, progressive defects in synaptic connectivity and ultrastructure occur alongside functional abnormalities in HCN channel activity and network dynamics, suggesting that pharmacological targeting of specific HCN channels could have significant therapeutic potential in human tauopathy.

## Results

### Increased tau phosphorylation and HCN channel expression in human post-mortem Alzheimer’s disease brain and in Tau35 mice

To examine a possible role for HCN channels in neurodegenerative pathology, we performed histological and biochemical analyses on hippocampal tissues from post-mortem AD and age-matched control human brain. Immunohistochemistry showed elevated HCN channels in the cornu ammonis (CA) 1 and CA3 hippocampal regions in AD brain, with statistically significant increases in the percentage of HCN1^+^ and HCN3^+^ neurons in CA3, and in HCN1^+^ neurons in the CA1 region (**Fig. 1, A and B**). Analysis of AD and control brain homogenates on western blots verified the increased HCN1 expression in AD hippocampus (**Fig. 1C**). Other synaptic markers, including synapsin-1, synaptotagmin and postsynaptic density protein-95, did not show significant differences between AD and control brain (**Fig. S1A**). As expected, phosphorylated tau (PHF1 epitope, Ser396/Ser404) and total tau were increased in AD compared to control brain (**Fig. 1D**).

**Figure 1.**
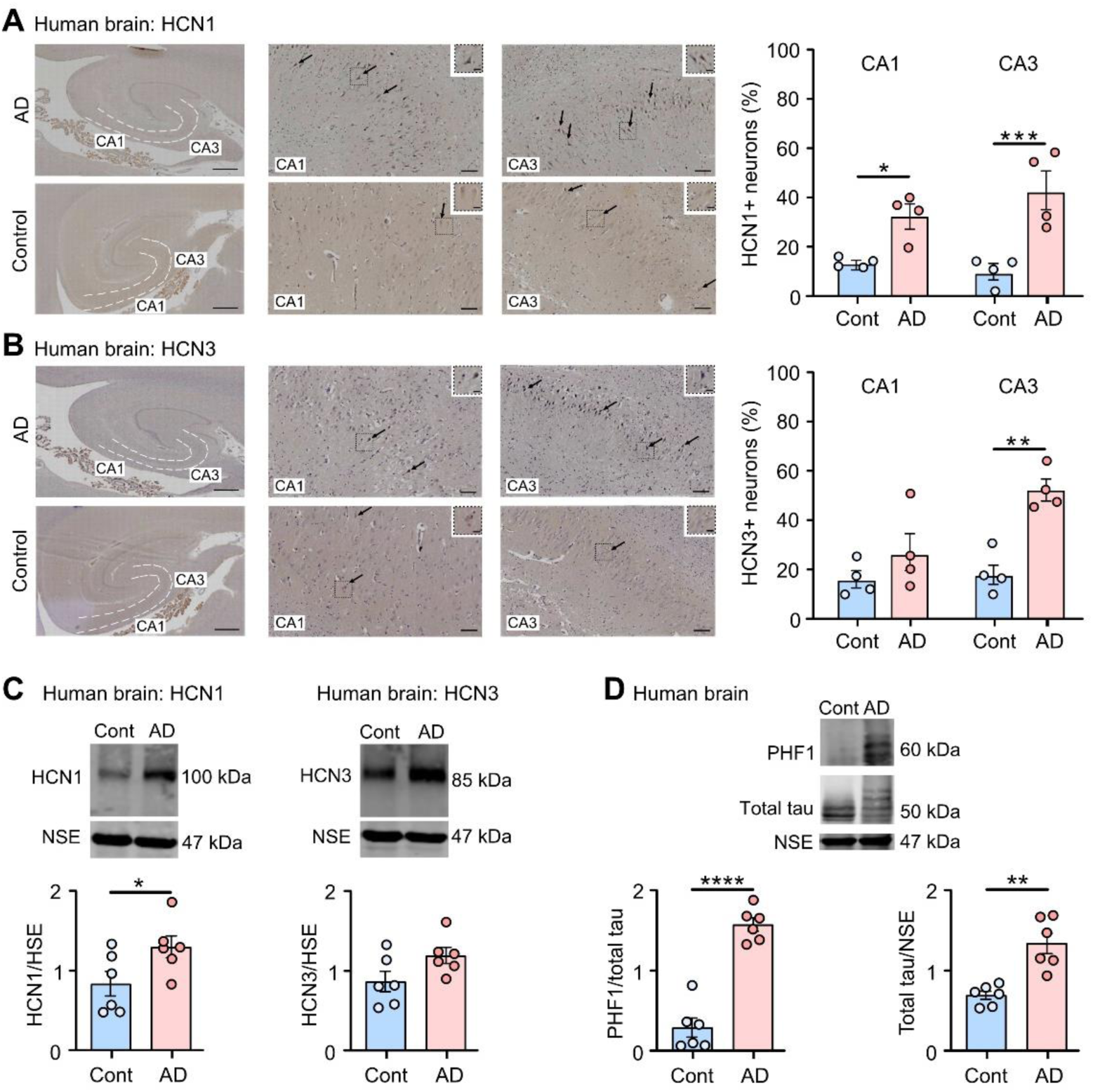
Increased HCN1 and HCN3 channel expression in human post-mortem Alzheimer’s disease brain. (**A,B**) HCN1 and HCN3 channel immunolabelling (arrows) in the hippocampus (cornu ammonis [CA] 1 and CA3 regions) of post-mortem Alzheimer’s disease (AD) and age-matched control (Cont) brain. Scale bars: left panels 1000 μm; center and right panels 100 μm; insets show higher magnification of indicated areas, scale bars: 20 μm. Quantification of the percentage of HCN^+^ neurons in the CA1 and CA3 regions is shown as mean ± SEM, n=4 brains per group. Student’s t-test, \**P* < 0.05, \*\**P* < 0.01, \*\*\**P* < 0.001. (**C,D**) Western blots of AD and control hippocampus probed with antibodies to HCN1, HCN3, phosphorylated tau (PHF1), total tau and neuron specific enolase (NSE). Quantification of the blots is shown as mean ± SEM, n=6 brains per group. Student’s t-test, \**P* < 0.05, \*\**P* < 0.01, \*\*\*\**P* < 0.0001.

Next, we examined the expression of HCN channels and synaptic markers in the brains of Tau35 mice, a model of progressive tauopathy that shows increased tau phosphorylation and deposition of abnormal tau species in parallel with deficits in cognitive and motor function, and reduced lifespan (*17*). Changes in HCN channels and synaptic markers as Tau35 mice age and disease develops, could have predictive value for determining whether network-wide structural and functional synaptic alterations are likely to progress and induce pathological alterations.

We first performed histological and biochemical analyses on hippocampus from post-symptomatic Tau35 (10 months old), when overt hippocampal tau pathology is apparent (*17*), and wild-type (WT) mouse brain, using antibodies to HCN1 and HCN3 (**Fig. 2**). We found statistically significant increases in the percentage of HCN3^+^, but not HCN1^+^, neurons in the CA3 region of the hippocampus in Tau35 mice (**Fig. 2, A and B**). Analysis of Tau35 and WT control brain homogenates on western blots showed significant increases in expression of both HCN1 and HCN3 channels, as well as increased tau phosphorylation in Tau35 mice (**Fig. 2, C and D**). In parallel, western blots of brain homogenates of pre-symptomatic mice (4 months old) were probed with antibodies to HCN1 and HCN3 to monitor whether changes in the expression of HCN channels occur before overt cognitive decline in Tau35 mice. Quantification of the blots showed significant increases in expression of HCN3, but not HCN1, together with increased phosphorylated tau in young Tau35 mice (**Fig. S2**). Marked decreases in the presynaptic marker synapsin-1, were observed in Tau35 mice at both ages, whereas synaptotagmin and postsynaptic density protein-95 were unchanged (**Fig. S2**). Since synapsin-1 is essential for clustering of synaptic vesicles (SVs) at the active zone (*18*), this reduction in synapsin-1 in Tau35 mice could affect SV organization and distribution at synapses.

**Figure 2.**
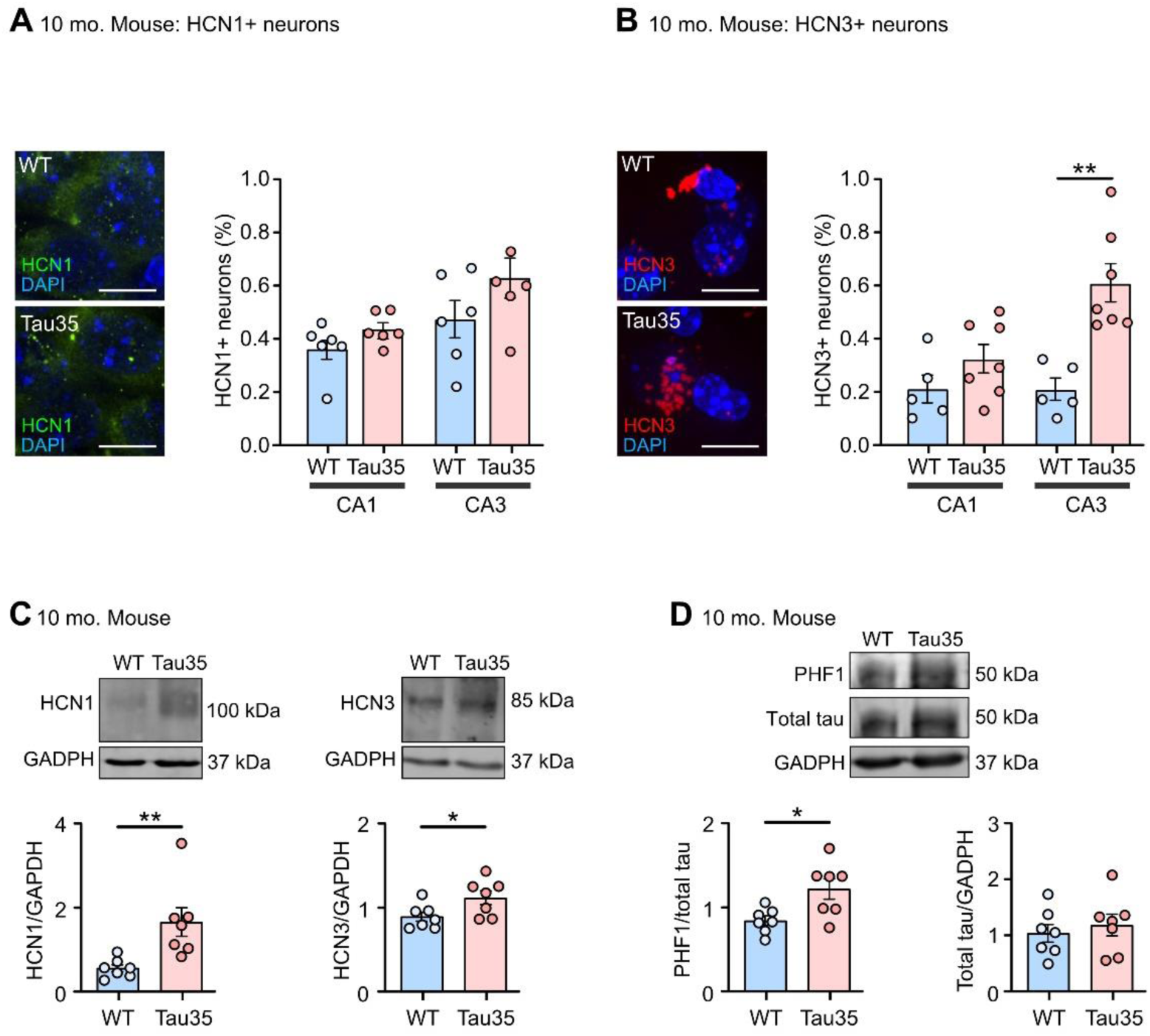
Increased HCN channel expression and tau phosphorylation in post-symptomatic Tau35 mouse brain. Immunofluorescence labeling of **(A)** HCN1 and **(B)** HCN3 in brain sections from wild-type (WT) and Tau35 post-symptomatic mice (aged 10 months). Scale bars: 10 μm. The percentage of HCN^+^ neurons in the CA1 and CA3 regions is shown in the graphs as mean ± SEM, n=5-7 brains per group. Student’s t-test, \*\**P* < 0.01. (**C,D**) Western blots of brain homogenates from WT and Tau35 mice aged 10 months, probed with antibodies to HCN1, HCN3, phosphorylated tau (PHF1), total tau and glyceraldehyde 3-phosphate dehydrogenase (GAPDH). Quantification of the blots is shown in the graphs as mean ± SEM, n=7 brains per group. Student’s t-test, \**P* < 0.05, \*\**P* < 0.01.

### Reduced dendritic branching and spine density in Tau35 mouse brain and primary hippocampal neurons

To look for morphological correlates of functional deficits in Tau35 neurons, we conducted Sholl analysis in Golgi-Cox stained hippocampus (CA1 region) in 10-month-old Tau35 and WT mice (**Fig. 3, A and B**). We found significant reductions in the number of dendritic branch points and mean dendrite length, but not in the soma area of Tau35 neurons compared to WT mice (**Fig. 3C**). Notably, expression of Tau35 significantly decreased overall arborization of basal dendrites of hippocampal neurons (two-way ANOVA, *P* < 0.0001), a possible correlate of reduced synaptic strength. In contrast, similar analysis of basal dendrites in 4-month-old mice, when Tau35 mice exhibit minimal tau pathology, did not reveal any significant differences in the number of dendritic branch points, mean dendrite length, or soma area (**Fig. S3, A and B**). Our results show that Tau35 expression reduces dendritic arborization and this loss of neuronal connectivity parallels the appearance of tau pathology and cognitive decline in Tau35 mice (*17*).

**Figure 3.**
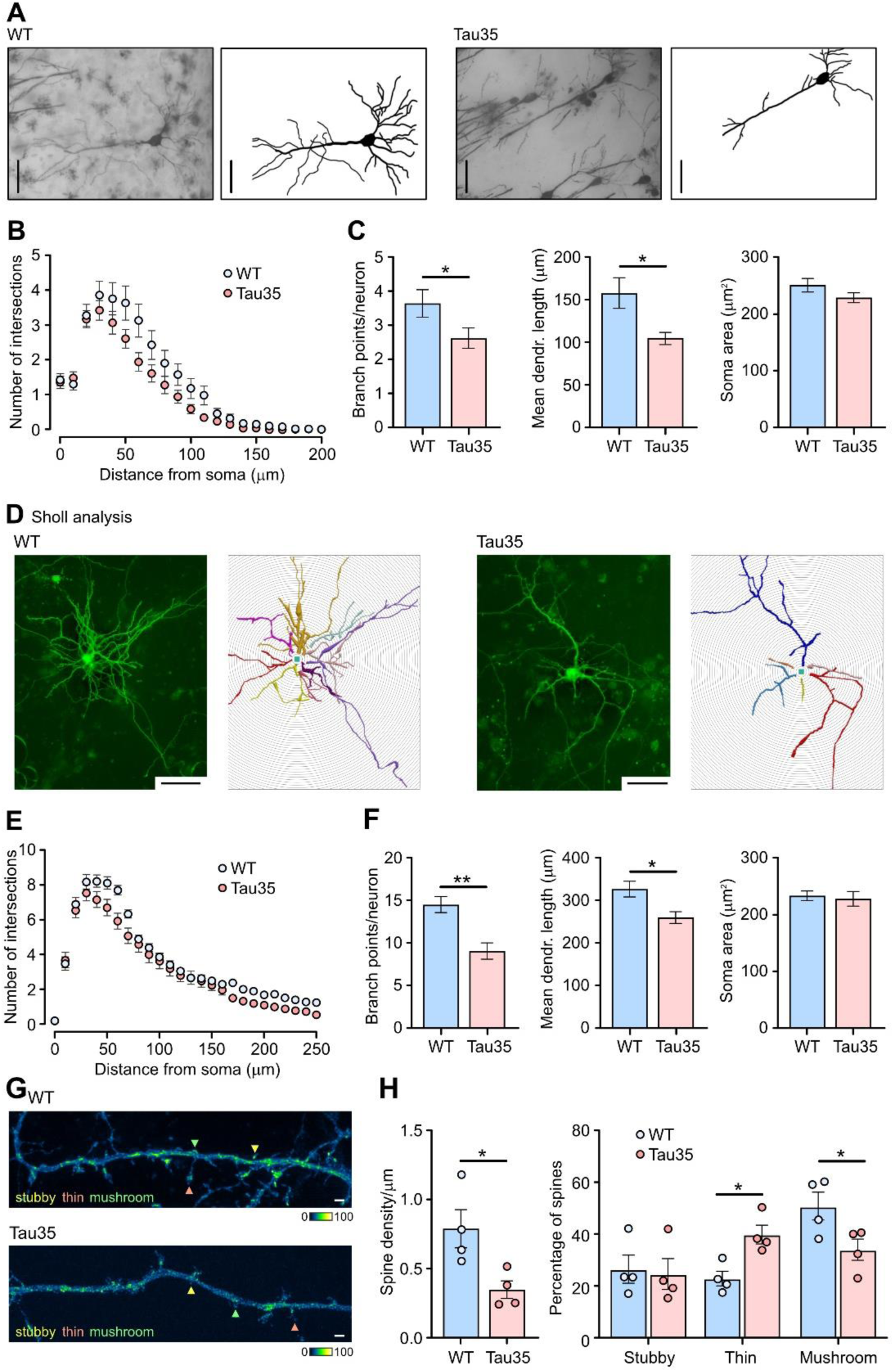
Progressive loss of dendritic complexity and spine density in Tau35 mouse brain and primary hippocampal neurons. (**A**) Golgi-Cox stained wild-type (WT) and Tau35 CA1 hippocampal neurons in 10-month-old mice. Scale bars: 50µm. (**B**) Sholl analysis demonstrates the reduced complexity of basal dendrites in Tau35 mice. Graph shows quantification of the mean ± SEM, n=40 neurons from 4-5 mice of each genotype. Two-way ANOVA, *P* < 0.001. (**C**) Graphs show the number of primary branch points per neuron, mean dendrite length, and soma area in CA1 neurons of WT and Tau35 mice, n=40 neurons from 4-5 mice of each genotype. Student’s t-test, * *P* < 0.05. (**D**) Fluorescence images (left) and Neurolucida drawings (right) of WT and Tau35 primary hippocampal neurons, transfected at 4 DIV with a plasmid expressing enhanced green fluorescent protein (eGFP) and imaged at 14 DIV. Scale bars: 50μm. (**E**) Sholl analysis of dendritic branching in WT and Tau35 hippocampal neurons at 14 DIV shows reduced dendritic complexity in Tau35 neurons. Graph shows quantification of the mean ± SEM, n=52 neurons of each genotype from 4-5 independent experiments. Two-way ANOVA, *P* < 0.001. (**F**) Graphs show the number of dendritic branch points, mean dendrite length, and soma area in WT and Tau35 hippocampal neurons at 14 DIV, n=52 neurons of each genotype, from 4-5 independent experiments. Student’s t-test, \**P* < 0.05, \*\**P* < 0.01. (**G**) Representative images of dendrites from WT and Tau35 hippocampal neurons transfected with a plasmid expressing eGFP and imaged at 14 DIV. Stubby spines (yellow), thin spines (orange) and mushroom spines (green) are indicated. Scale bars, 1 mm. (**H**) Graphs show quantifications of spine density (number of spines per μm dendrite length) and percentage of each spine type in WT and Tau35 hippocampal neurons at 14 DIV; n=50 neurons of each genotype from 4 independent experiments. Student’s t-test, \**P* < 0.05.

We established an *in vitro* model of Tau35 neurons and conducted a Sholl analysis in enhanced green fluorescent protein (eGFP)-expressing hippocampal neurons, which were transfected at 4 DIV and imaged at 6, 9, and 14 DIV. In 6 DIV neurons, we observed no effect of Tau35 on dendritic branching, mean dendrite length or soma area (**Fig. S3C**), whilst there was a modest, statistically significant reduction in branching by 9 DIV (**Fig. S3D**, two-way ANOVA, *P* < 0.01). However, there was a striking reduction in the dendritic complexity of Tau35 neurons at 14 DIV (**Fig. 3, D and E**, two-way ANOVA, *P* < 0.001). Specifically, the number of dendritic branch points in Tau35 neurons decreased by 44% and the mean dendrite length reduced by 20% compared to WT neurons, without any change in soma area (**Fig. 3F**). Correct formation of the dendritic arbor has important consequences for synaptic function in neurons. As such, this marked reduction in dendritic complexity suggests that the presence of Tau35 could result in decreased synaptic activity and/or synaptic strength.

Next, to provide further insights into functional changes, we characterised dendritic spine morphology, given that both spine number and structure are known correlates of synaptic efficacy (*19, 20*). The number of each morphologically defined dendritic spine type was determined from three-dimensional reconstructions of eGFP-expressing hippocampal neurons at 14 DIV (**Fig. 3G**). Dendritic spine density was reduced by 59% (*P* < 0.05) in Tau35 neurons (**Fig. 3H**). Moreover, Tau35 expression specifically reduced the proportions of stubby and mushroom spines (**Fig. 3H**, *P* < 0.05). This dramatic decrease in spine number suggests that Tau35 expression has a potentially negative impact on synaptic transmission and could limit synaptic plasticity (*21*) and enhance synaptotoxicity.

### Ultrastructural changes in hippocampal presynaptic terminals of Tau35 mice

We next used an ultrastructural approach to characterize the possible impact of Tau35 on presynaptic structure in the hippocampal CA1 region in mice (**Fig. 4A**). In representative electron micrograph sections taken from mice aged 4 months we recorded a significant reduction (22%) in the number of synaptic vesicles per terminal in Tau35 versus WT hippocampus (**Fig. 4B**). In Tau35 mice aged 10 months, this difference was even more pronounced (39% reduction in Tau35 mice, **Fig. 4B**). In contrast, there were no significant changes in the presynaptic area, the diameter of synaptic vesicles, or the percentage of docked synaptic vesicles in Tau35 mice at either age (**Fig. S4, A to C**). These findings indicate that Tau35 has a marked and sustained negative impact on correlates of presynaptic strength that progress further with aging.

**Figure 4.**
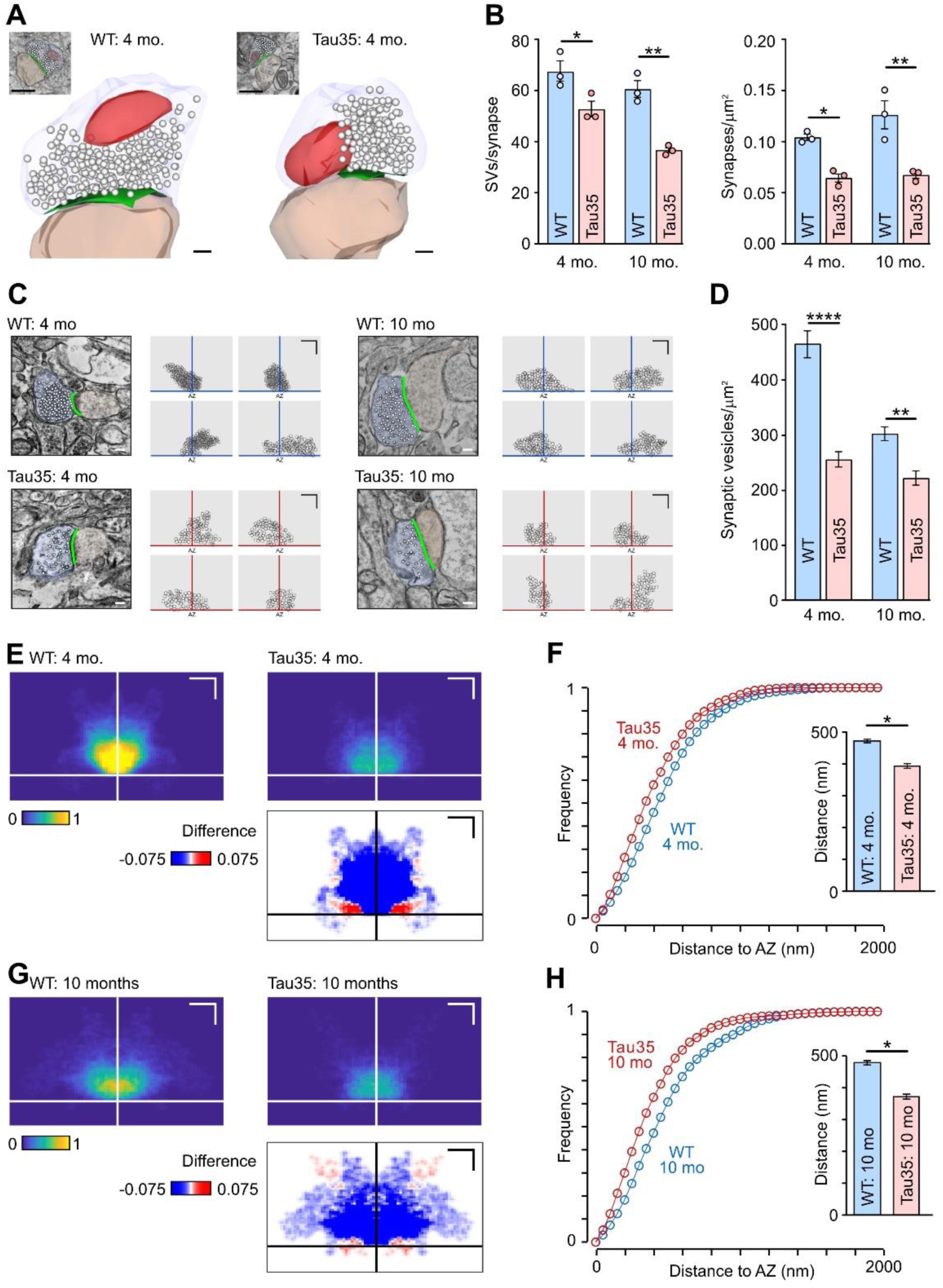
Ultrastructural analysis of synapses in wild-type and Tau35 mouse brain. (**A**) Representative electron micrographs (top, inset) and 3D serial reconstructions of wild-type (WT) and Tau35 CA1 hippocampal synapses in mice aged 4 months. Presynaptic (light blue) and postsynaptic (pink) structures are separated by the electron-dense active zone (green), which is surrounded by synaptic vesicles (white). Mitochondria are shown in red. Scale bars: 500 nm. (**B**) Plots showing the number of synaptic vesicles (SVs) per synapse **(left)** and synaptic density **(right)** of the CA1 region of the hippocampus in WT and Tau35 mice aged 4 and 10 months. Two-way ANOVA, n=100-150 synapses from 3 mice of each genotype, \**P* < 0.05, \*\**P* < 0.01. (**C**) Representative electron micrographs of CA1 synapses in WT and Tau35 mice aged 4 months (left) and 10 months (right) from randomly selected samples used for analysis of spatial organization. Presynaptic (light blue) and postsynaptic (pink) structures, and synaptic vesicles (white) are pseudocoloured. Scale bars: 100 nm. Schematics showing representative vesicle clusters in synaptic terminals in WT and Tau35 mice are shown on the right. Circles indicate individual synaptic vesicles; orange and blue crosshairs indicate active zone centers. Scale bars: 400 nm. (**D**) Plot showing the packing density of synaptic vesicles for each condition. Two-way ANOVA, \*\**P* < 0.01, \*\*\*\**P* < 0.0001. **(E-H)** Color-coded spatial frequency density plots show mean vesicle density distributions for **(E)** WT and **(G)** Tau35 mice (4 and 10 months old), n=20 synapses. Crosshairs indicate active zone centers. Scale bars: 200 nm. Student’s t-test, \**P* < 0.05. Lower panels in **E and G** show difference images of signal predominating in WT (blue) and Tau35 (red) clusters. Cumulative frequency distance plots **(F and H)** show the distribution of synaptic vesicles relative to the nearest point on the active zone.

To examine possible alterations in the spatial organization of synaptic vesicles between conditions, we mapped vesicle positions in a randomly selected sample of representative middle sections, which included the active zone. Strikingly, the packing density of synaptic vesicles was lower in Tau35 mice at both age points, a finding confirmed by quantifying the number of vesicles per unit area using concave hulls to define cluster boundaries (**Fig. 4, C and D**). This implies that Tau35 could be affecting the molecular machinery that serves to cluster and corral vesicles at terminals. Next, we characterized vesicle positions within the terminal architecture with respect to the active zone center. These coordinates were used to generate color-coded mean spatial frequency distribution maps, which revealed that Tau35 synapses had a lower density vesicle cloud close to the active zone than WT synapses at the same age (**Fig. 4, E-H**). We quantified this change in cluster organization by measuring the distance from each vesicle to its nearest point on the active zone, showing that cumulative frequency distance plots were left shifted in Tau35 terminals versus

WT at both time points and this yielded shorter average distances (**Fig. 3, F and H**), presumably reflecting the smaller cluster sizes overall. Strikingly, difference plots for the Tau35 versus WT distribution maps show that vesicles in Tau35 synapses favor aggregation at sites adjacent to the active zone and at the lateral edges of the cluster (**Fig. 4, E-G**, lower panels). Since these are likely regions for vesicle retrieval (*22, 23*), it suggests a possible nanoscale correlate of recycling deficits at Tau35 terminals.

In addition to synaptic vesicle-driven fast transmission, slower synaptic signaling occurs via neuropeptides packaged in dense-core vesicles (DCVs). Therefore, we quantified the percentage of synapses harboring DCVs in Tau35 and WT mice. At 4 months of age, there was no difference between the genotypes (**Fig. S4D**). However, 10-month-old Tau35 mice exhibited a marked (74%) reduction in the percentage of hippocampal synapses with DCVs (**Fig. S4D**). This reduction in DCV-containing synapses in Tau35 mice occurs at a time when overt cognitive dysfunction and tauopathy are apparent in these animals, suggesting that the lack of synaptic DCVs could result in compromised release of neuropeptides and thereby further inhibit information transfer in neural networks.

### Tau35 influences I_h_-dependent sag and excitatory postsynaptic current kinetics in hippocampal neurons

To look for functional correlates of morphological deficits in Tau35 neurons, we performed a detailed electrophysiological analysis using hippocampal cultures. First, to confirm the validity of this *in vitro* preparation, we used immunofluorescence labeling to show that all four HCN channels are expressed at 14 DIV (**Fig. 5A, Fig. S5A**). Strikingly, the same general pattern of increased HCN channel expression was seen in these cultured neurons as we observed in human AD and in Tau35 mice (**Figs. 1 and 2**). Specifically, there were significant increases in both HCN1 (59%) and HCN3 (38%) in Tau35 neurons versus WT (**Fig. 5B**), but no differences in HCN2 or HCN4 channel expression (**Fig. S5A**). Likewise, quantification of HCN1 and HCN3 channels on western blots showed increases of 52% and 61% respectively, in Tau35 neurons (**Fig. 5C**). PHF1 labeling was also increased more than two-fold in Tau35 hippocampal neurons at 14 DIV, indicating that tau phosphorylation in Tau35 mice is elevated from an early stage of neuronal development (**Fig. S5B**). Our *in vitro* model therefore recapitulates the key features of human AD and Tau35 mice and is thus a disease-relevant tool in which to assess functional changes.

**Figure 5.**
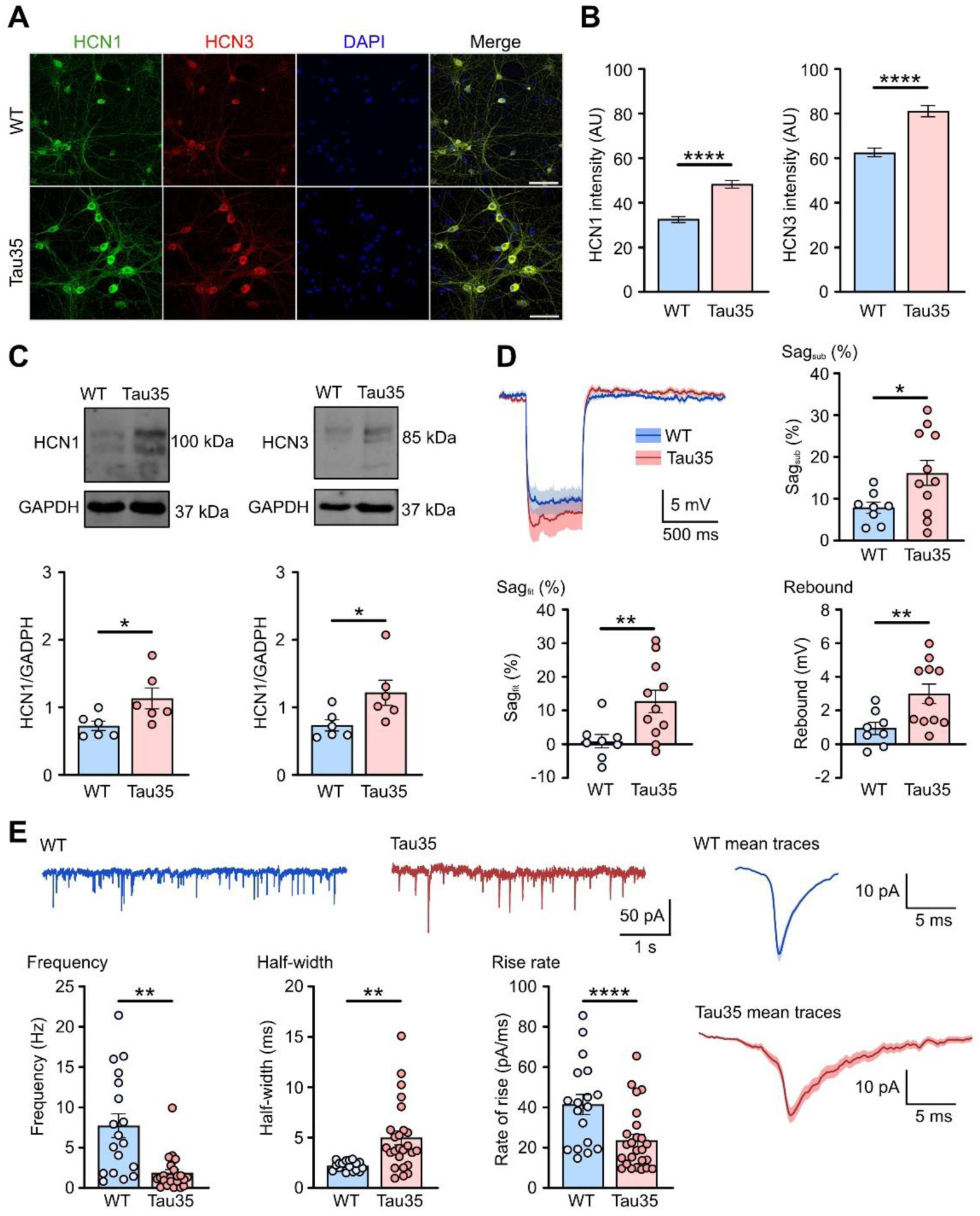
Increased HCN channel expression and functional changes in Tau35 hippocampal neurons. (**A**) Immunofluorescence labeling of wild-type (WT) and Tau35 mouse hippocampal neurons at 14 DIV with HCN1 and HCN3 antibodies, and DAPI staining of nuclei. Scale bars: 50 μm. (**B**) Graphs show quantification of mean fluorescence intensity (± SEM) of HCN1 and HCN3. n=150 (WT) and n=120 (Tau35) neurons, from 3 independent experiments. Student’s t-test, \*\*\*\**P* < 0.0001. (**C**) Western blots of lysates of primary hippocampal neurons (14 DIV) from WT and Tau35 mice, probed with antibodies to HCN1 (left) or HCN3 (right), and glyceraldehyde 3-phosphate dehydrogenase (GAPDH). Quantification of the blots is shown in the graphs as mean ± SEM; n=6 independent experiments. Student’s t-test, \**P* < 0.05. (**D**) Average traces ± SEM boundaries of V_m_ hyperpolarization by injection of −100 pA, 500 ms current in WT and Tau35 mouse hippocampal neurons (11-16 DIV). Graphs show sag_sub_, sag_fit_, and rebound potential. Graphs show quantification of the mean ± SEM; n=8 (WT) and n=11 (Tau35) neurons. Student’s t-test, \**P* < 0.05, \*\**P* < 0.01. (**E**) Spontaneous excitatory postsynaptic currents (sEPSCs) in WT and Tau35 hippocampal neurons (11-16 DIV). Left panel: representative traces; right panel: average traces ± SEM. Graphs show frequency, half-width, and rate of rise of sEPSCs. Graphs show quantification of the mean ± SEM; n=18 (WT) and n=25 (Tau35) neurons. Student’s t-test, \*\**P* < 0.01, \*\*\*\**P* < 0.0001.

Next, to establish Tau35 effects on neuronal excitability in early development, we performed whole-cell recordings from cultured hippocampal neurons at 11-16 DIV. Specifically, we recorded the deflection of the plasma membrane voltage (V_m_) in response to a hyperpolarizing current step to explore the I_h_-dependent sag and rebound (*24*) properties in Tau35 and WT neurons (**Fig. 5D**). Sag was quantified by the subtraction of the negative peak (sag_sub_) and of the extrapolated first exponential fit of the V_m_ decay (sag_fit_) upon the injection of a hyperpolarizing step (*24*) (illustrated in **Fig. S5C**). Sag_sub_ increased approximately two-fold and sag_fit_ was elevated almost 15-fold in Tau35 versus WT hippocampal neurons. The increased sag was accompanied by a three-fold rise in rebound potential in Tau35 neurons when the hyperpolarizing current step ended (**Fig. 5D**). Since sag potentials increase in response to activation of I_h_ current through HCN channels (*25*), our data strongly suggest that the augmented sag in Tau35 neurons is due to increased expression of HCN1 and HCN3 channels (**Fig. 5, A-C**). Notably, we detected no changes in the biophysical properties of the currents generated by non-inactivating potassium channels, including the maximal conductance (G_max_) and the half-activation voltage (V_1/2_) in Tau35 neurons (**Fig. S5, D and E**).

To investigate whether the changes that we attribute to HCN channels affected downstream information signaling, we next recorded spontaneous excitatory postsynaptic currents (sEPSCs) in Tau35 and WT neurons at 11-16 DIV (**Fig. 5E**). We found that Tau35 neurons exhibited a marked reduction (77%) in their sEPSC frequency (**Fig. 5E**), without changes in amplitude (**Fig. S5F**), consistent with Tau35 reducing the probability of presynaptic vesicle release. The half-width of sEPSCs in Tau35 neurons increased more than two-fold, and the rate of rise reduced to 43% compared to WT neurons (**Fig. 5E**), which may contribute to temporally less precise spike responses (*26*). Given the established link between asynchronous vesicle release and the timing of sEPSCs (*27, 28*) we propose that these Tau35-associated changes in EPSC kinetics might reflect changes in synaptic vesicle recycling.

Finally, to assess the effect of Tau35 on the electrogenic properties of the plasma membrane, we evoked action potentials using a 300 pA, 500 ms square current injection and measured their waveform properties (**Fig. S5F**). None of the action potential properties measured, including peak, threshold, maximum rate of rise, peak width, EPSC amplitude, time constant (tau), or rate of decay were affected by Tau35 (**Fig. S5F** and data not shown).

## Discussion

Proper maintenance of synaptic structure and function is critical for cognitive processing, and synaptic dysfunction is an early correlate of dementia and related tauopathies (*29*). However, the molecular mechanisms that underlie the synaptic damage caused by tau deposition are not understood. Here we demonstrate pathological alterations in the expression and localization of selected HCN channels in human AD and Tau35 mouse brain that provide a basis for explaining wide-ranging deficits in neuronal signaling and connectivity (**Fig. 6**).

**Figure 6.**
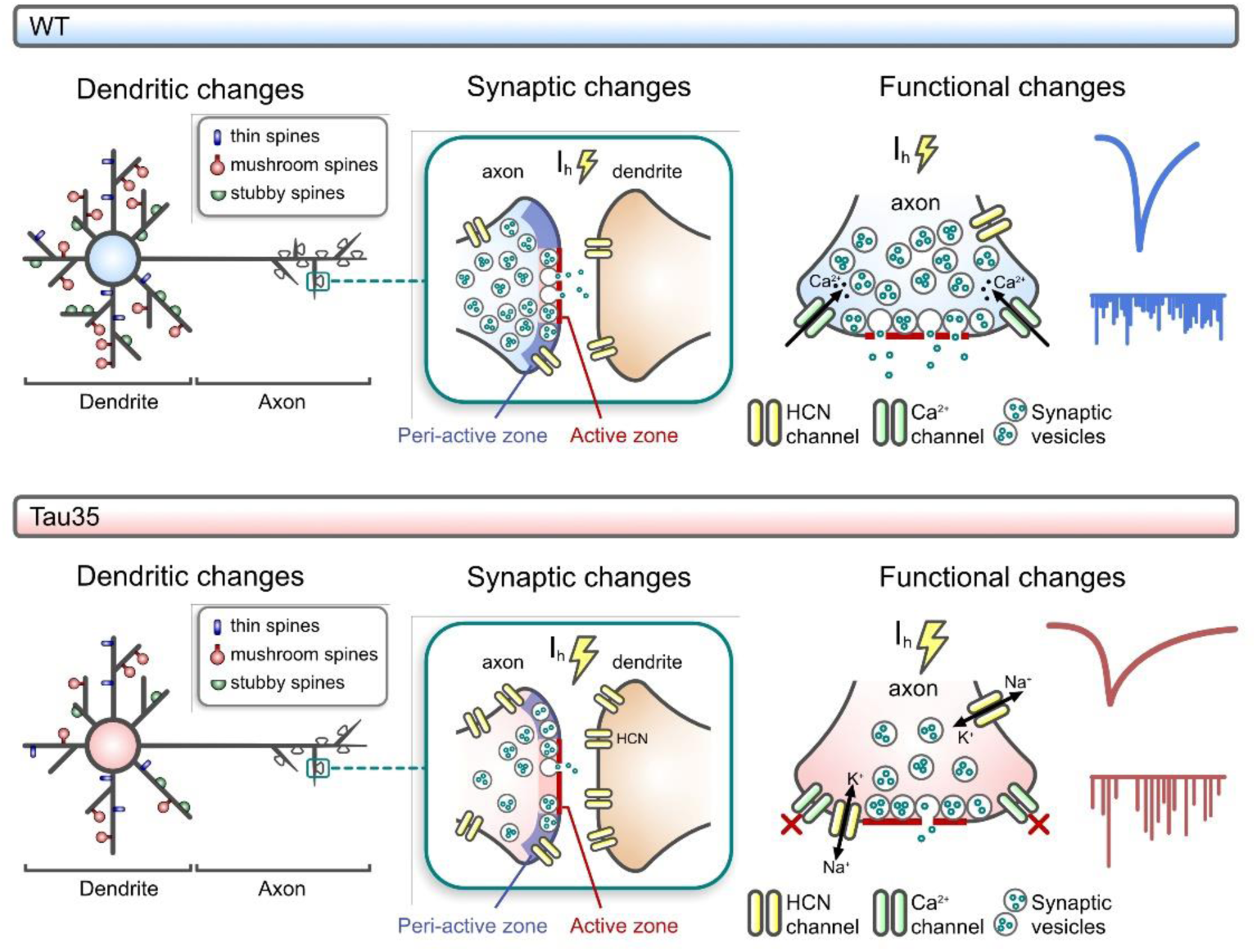
Proposed model of Tau35-induced HCN channelopathy and synaptic dysfunction. Expression of Tau35 results in reduced dendritic complexity and spine density, accompanied by alterations at synapses, including decreased synaptic vesicle density and clustering at peri-active zones. Increased expression of hyperpolarization-activated cyclic nucleotide-gated (HCN) channels is paralleled by increases in I_h_-dependent sag voltage and half-width of spontaneous excitatory postsynaptic currents (sEPSCs), along with reductions in the frequency and rate of rise of sEPSCs. WT: wild-type.

HCN channels regulate a range of cellular properties, including membrane resistance, intrinsic membrane excitability and synaptic integration (*6*), and are emerging as key players in the pathogenesis of several neurodegenerative diseases (*13*). Expression of HCN channels determines I_h_ current generation and fine tunes communication between hippocampal subregions and cortical or subcortical networks (*30, 31*). As HCN channels regulate hippocampus-dependent learning and memory (*32*), the imbalance of HCN channel expression in AD and Tau35 brain is a likely candidate for driving abnormal communication between hippocampal and cortical structures that triggers changes in memory formation and retrieval.

In Tau35 mice (*17*), progressive increases in HCN1 and HCN3 ion channel expression are accompanied by an array of dendritic, synaptic and ultrastructural changes as well as the development of tau pathology, with increases in dendritic branching observed only during advanced tauopathy. We note that expression of P301L mutant tau in rTg4510 transgenic mice results in a similar, robust reduction in dendritic arborization at late stages of disease (*15*). In the same tau transgenic mouse, structural and functional synaptic alterations were observed in the early stages of tauopathy both *in vivo* (*33*) and *in vitro* (*34*). Our findings have important mechanistic implications for disease because the degree of dendritic branching influences the generation of new synapses, neuronal function, and cognitive performance, all of which are impaired in tauopathies. Dendrites are also a major source of neuropeptides and their release, even independently of electrical activity, enables sustained re-organization of neuronal networks (*35*). Our observation of a marked reduction in synapses harboring DCVs in the hippocampus of aged Tau35 mice, also mirrors the reduction in hippocampal DCV peptide density reported in AD (*36*).

Recent studies have shown that pathogenic tau alterations in axons precede disease-associated changes in the somatodendritic compartment (*37*). Indeed, in the hippocampus of young pre-symptomatic Tau35 mice, we found enhanced phosphorylated tau load as well as marked reductions in synapse density and in the number of synaptic vesicles in synaptic terminals, changes that precede the appearance of overt tau pathology in these animals (*17*). We also observed altered presynaptic cytoarchitecture in the brains of young Tau35 mice, with more loosely distributed synaptic vesicles without a prominent core cluster, at sites adjacent to the active zone and at the lateral edges of the cluster. The reduction in synapsin, a key synaptic vesicle tether (*38, 39*), in Tau35 neurons could potentially explain this loosened cluster phenotype. Indeed, consistent with our observations, synapsin knockout mice show similar reductions in vesicle number and vesicle density in hippocampal terminals and increased clustering of vesicles in peri-synaptic regions (*40*). Vesicle organization at presynaptic terminals of hippocampal synapses has been linked to synaptic efficacy (*41–43*), suggesting that disruptions in the integrity of synaptic vesicle clusters could underlie deficits in synaptic signaling in Tau35 mice.

Changes in expression of voltage-gated channels encoded by the HCN1-4 gene family have been reported in multiple neurological disorders (*13*). These channels have major roles in controlling synaptic transmission, integration of synaptic input and neuronal excitability (*44*). In particular, HCN1 has been shown to mis-localize and become dysfunctional in models of dementia (*14, 34*). Impaired trafficking of HCN1 channels to distal dendrites or increased HCN expression (*14, 45*) is associated with increased I_h_-dependent sag potential in multiple transgenic models of AD (*14–16*), suggesting a possible role for HCN in the development of disease.

We used cultured Tau35 hippocampal neurons to better understand how changes in the expression of HCN channels may drive dynamic structural and functional synaptic changes. We first examined dendritic spine morphology in cultured Tau35 hippocampal neurons. This is important because activity-dependent spine remodeling is critical for neuronal communication (*46*) and synaptic strength is regulated by the number and size of individual dendritic spines (*19, 20*). In Tau35 hippocampal neurons we observed reduced density of dendritic spines and loss of mature spines, which adversely affects synaptic strength and further supports the view that tau truncation compromises synaptic communication.

Reduced dendritic and synaptic complexity have been linked to altered resonance in hippocampal pyramidal neurons of rTg4510 mice overexpressing P301L tau (*15, 34*). Similarly, we reported changes in resonance in Tau35 mice, including reduced impedance in older animals upon injection of an oscillating current (*47*). Here we observed altered membrane dynamics in Tau35 primary hippocampal neurons and identified increased voltage-sensitivity of non-inactivating potassium channels as a possible cause of the decreased impedance and voltage-dependent input resistance caused by Tau35 expression. Conversely, Tau35 hippocampal neurons did not exhibit changes in either input resistance or biophysical properties of voltage-gated potassium channels that generate M-currents (I_M_) (*48*). However, we did detect increased I_h_-dependent sag of the V_m_ deflection upon injection of negative current, paralleled by elevations in the I_h_-generating HCN1 and HCN3 channels. Interestingly, both I_h_ and I_M_ currents play critical roles in regulating plasma membrane excitability in the presence of oscillating potentials, driving neuronal resonance (*49*). The interplay of I_h_ and I_M_ has recently been reported to regulate intrinsic neuronal excitability and to affect disease progression in a mouse model of amyotrophic lateral sclerosis (*50*). Neurons with strong I_h_ currents are good resonators and operate as bandwidth filters (*51, 52*). In contrast, inhibited I_h_ and increased I_M_ current results in low resonance, causing the neuron to function as a low-pass filter and attenuating higher frequencies (*53*). Our data demonstrate that Tau35 is associated with increased I_h_-related events in primary hippocampal neurons whereas, in aged animals, Tau35 is associated with increased I_M_ (*47*). Notably, any alteration to the I_h_/I_M_ ratio caused by Tau35 expression could interfere with intrinsic neuronal excitability and the ability of hippocampal neurons to respond to specific stimuli, further disrupting neuronal connectivity.

Tau35 primary hippocampal neurons develop sEPSCs that were less frequent, wider, and with slower rise times than those of WT neurons. These currents reflect alterations in presynaptic glutamate release and subsequent activation of post-synaptic AMPA and NMDA receptors (*54*). Notably, fluctuation in the release of presynaptic vesicles has been reported to profoundly affect both the timing of sEPSCs (*27, 28*) and synaptic vesicle distribution along the active zone (*55*). The changes we observed in sEPSCs in Tau35 hippocampal neurons could, therefore, be caused by asynchronous release of presynaptic vesicles.

In conclusion, this study identifies previously unreported roles for HCN1 and HCN3 channels in the development and progression of tauopathy (**Fig. 6**). Our results demonstrate that increased expression of HCN channels is likely responsible for the increased sag voltage apparent in Tau35 neurons. The aberrant signaling induced by Tau35 coincides with changes in synaptic ultrastructure, reducing the density of synaptic terminals as well as the clustering of synaptic vesicles in individual terminals, adversely affecting network connectivity. Importantly, Tau35 also disrupts synaptic vesicle positioning in synaptic terminals, resulting in excessive accumulation of synaptic vesicles in the peri-active zone. Our results suggest that selective targeting of specific HCN channels with pharmacological agents could potentially protect synapses and prevent disease progression in human tauopathy.

## Materials and Methods

### Human tissue

#### Ethics Statement

All cases were neuropathologically assessed in accordance with standard criteria following consent for autopsy and research. All studies were conducted under the ethical approval of the King’s College London and MRC London Neurodegenerative Diseases Brain Bank.

#### Preparation of human brain homogenates for western blots

Frozen human hippocampi from AD and age-matched controls were obtained from the London Neurodegenerative Diseases Brain Bank (King’s College London, UK). Tissue was homogenized in 10 volumes of ice-cold radio-immunoprecipitation assay (RIPA) buffer (150 mM NaCl, 1 mM ethylenediaminetetraacetic acid [EDTA], 50mM Tris-HCl, 1% [v/v] NP-40, 0.5% [w/v] sodium deoxycholate, 0.1% [w/v] sodium dodecyl sulfate [SDS]), supplemented with protease inhibitor cocktail (cOmplete^™^, EDTA-free, Merck Millipore) and phosphatase inhibitor (PhosSTOP, Sigma-Aldrich) using a glass Dounce homogenizer. Hippocampal homogenates were centrifuged at 10,000g for 15 min at 4°C and the supernatants were stored at −80°C.

#### Immunohistochemistry of human brain tissue

Human brain sections of 7 μm thickness were cut from formalin fixed paraffin-embedded tissue blocks and deparaffinized in xylene. Endogenous peroxidase was blocked by immersion in methanol with H_2_O_2_ for 30 min and antigen retrieval was carried out using 20 min microwave heating in citrate buffer, pH 6. After blocking in normal serum, primary antibody was applied overnight at 4°C. After two washes in Tris-buffered saline (TBS), sections were incubated with biotinylated secondary antibody (DAKO), followed by avidin/biotinylated enzyme complex (Vectastain Elite ABC kit, Vector Laboratories). Finally, sections were incubated for 15 min with 3,3′-diaminobenzidine chromogen (0.5 mg/ml) (Sigma-Aldrich) in Tris-buffered saline (pH 7.6) containing 0.05% H_2_O_2_. Sections were counterstained with Harris’s haematoxylin, and images acquired using a VS120EL100W, Virtual Slide Microscope (Olympus). Automated image analysis and quantification were performed on antibody-labelled human hippocampal sections using customized digital analysis APPs (Visiopharm A/S).

### Animals

#### Ethics Statement

Tau35 mice were generated by targeted knock-in to the *Hprt* locus under the control of the human tau promoter as described previously (*17*). All procedures were conducted in accordance with the Animals (Scientific Procedures) Act, 1986, following approval by the local ethical review committee. All procedures conformed to the ARRIVE guidelines 2.0.

#### Preparation of mouse brain homogenates for western blots

Mice were sacrificed by cervical dislocation. Brains were removed and snap-frozen in dry-ice and stored at −80°C. Tissue was lysed by ultrasonication (sonication parameters: amplitude 40%; pulse 4 s; time: 30 s) using a Vibra-Cell ultrasonic liquid processor model no. VCX 130 (Sonics and Materials, Newton, CT, USA) in ice-cold RIPA buffer (150 mM NaCl, 1 mM ethylenediaminetetraacetic acid [EDTA], 50mM Tris-HCl, 1% [v/v] NP-40, 0.5% [w/v] sodium deoxycholate, 0.1% [w/v] sodium dodecyl sulfate [SDS]), supplemented with protease inhibitor cocktail (cOmplete^™^, EDTA-free, Merck Millipore) and phosphatase inhibitor (PhosSTOP, Sigma-Aldrich), followed by incubation on ice for 3 min, repeating this cycle three times. Lysed tissue was centrifuged at 10,000g for 15 min at 4°C and the supernatants were stored at −80°C.

#### Preparation of mouse brain tissue for immunolabeling and Golgi-Cox staining

Mice were sacrificed using terminal anaesthesia, perfused with phosphate-buffered saline (PBS) and post-fixed in 4% (w/v) paraformaldehyde (PFA) overnight at 4°C. Brain sections (50-200 µm) were prepared using a VT1000 S Vibrating blade microtome (Leica Biosystems) and stored free-floating in cryoprotectant (30% (v/v) ethylene glycol, 15% (w/v) sucrose in PBS) at −20°C.

For immunofluorescence, 50 µm sections were washed in PBS, blocked in 3% (v/v) goat serum (Sigma-Aldrich), in 0.1% (v/v) Triton X-100 (Thermo Scientific) in PBS, and incubated in primary antibody (Table 2) in PBS overnight at 4°C. After washing, sections were incubated in the appropriate fluorophore-conjugated secondary antibody (Table 2) for 4-6 h at 4°C, and counterstained with 4′,6-diamidino-2-phenylindole (DAPI), before mounting in fluorescence mounting medium (S3023, Agilent Dako). Whole brain sections were imaged using a VS120EL100W, Virtual Slide Microscope (Olympus) equipped with x20, 0.75 NA and x40, 0.95 NA, air objectives. Automated image analysis and quantification were performed on antibody-labeled mouse brain sections using customized digital analysis APPs (Visiopharm A/S). High magnification images of neurons in the CA1 and CA3 regions of the hippocampus were collected using the Nikon Eclipse Ti Inverted Spinning Disk Confocal System (100x oil objective, automated acquisition using the NIS-Elements JOBS module).

For Golgi-Cox staining, 200 μm sections were stained according to the manufacturer’s instructions (sliceGolgi Kit, Bioenno Lifesciences) and imaged using a Nikon Ti-E two-camera microscope (×60, 1.4 NA, oil objective). Three-dimensional digital reconstruction of dendritic, axonal and spine structures was performed using Neurolucida software (BFM Bioscience). Whole brain sections were imaged using a VS120EL100W, Virtual Slide Microscope (Olympus) equipped with x20, 0.75 NA and x40, 0.95 NA, air objectives.

#### Preparation of mouse brain for ultrastructural analysis

Mice were sacrificed using terminal anaesthesia, perfused with PBS and post-fixed in 4% (w/v) PFA, 0.1% (w/v) glutaraldehyde in PBS, overnight at 4°C. Brains were removed and 300 μm transverse brain sections were prepared using a vibratome (VT1000 S Vibrating blade microtome, Leica Biosystems). Fixative was replaced with 0.2 M sodium cacodylate and 0.2 M sodium cacodylate, supplemented with 0.02 M CaCl_2_. Sections were treated with 2% (w/v) osmium tetroxide for 1 h on ice. After washing in ultra-pure water, sections were stained for 20 min in 0.1% (w/v) thiocarbohydrazide at ambient temperature, followed by three washes in water. Sections were treated with 2% (w/v) osmium tetroxide in 0.2 M sodium cacodylate, supplemented with 0.02 M CaCl_2_ and 1.5% (w/v) potassium ferrocyanide. After a further water wash, sections were stained by incubating in 1% (w/v) uranyl acetate overnight at 4°C. Stained sections were dehydrated in ascending concentrations of ethanol, then in acetone, before flat embedding in ascending concentrations of Durcupan resin (25% [v/v] for 2 h, 50% [v/v] for 2 h, 70% [v/v] for 2 h, 100% [v/v] overnight). Fresh resin was added and polymerised using a UV light translinker for 3-4 days at 4°C (UVP TL-2000 Translinker). The stained and embedded hippocampus was dissected, glued onto a BEEM capsule, mounted in an ultramicrotome (Leica), and ultrathin 70 nm sections were collected on hexagonal 300-mesh nickel grids (3.05 mm; Agar Scientific).

### Western blots

Protein concentration was determined using a bicinchoninic acid protein assay, according to the manufacturer’s instructions (Pierce™ BCA Protein Assay Kit, Thermo Fisher Scientific). Samples in Laemmli sample buffer were incubated at 95°C for 10 min and electrophoresed on 10% (w/v) SDS-polyacrylamide gels. Separated proteins were transferred to polyvinylidene difluoride (PVDF) membranes, blocked in Intercept^®^ (TBS) Blocking Buffer (LI-COR Biosciences) and incubated in primary antibodies (Table 1), overnight at 4°C. Following washing in TBS containing 0.02% (v/v) Tween 20, membranes were incubated with appropriate fluorophore-conjugated secondary antibodies (Table 1) for antigen detection and imaged (Odyssey® imager, LI-COR Biosciences). ImageStudio^TM^ Lite software (LI-COR Biosciences) was used for quantification of western blots.

### Cell culture

Primary hippocampal neurons were prepared from embryonic (E) day 16.5-18.5 Tau35 and WT mice (*17*). and cultured as described previously (*56*). Neurons were transfected (Lipofectamine 2000, Thermo Fisher Scientific) with a plasmid expressing eGFP at 4 DIV and fixed at 6, 9, and 14 DIV to monitor dendritic branching and quantify spine density. Untransfected neurons were cultured for 6, 9, and 14 DIV and either fixed in 4% (w/v) paraformaldehyde (PFA) for 15min and labelled with antibodies for immunofluorescence or lysed for analysis on western blots. For immunofluorescence staining, non-specific sites were blocked for 30 min in 3% (v/v) goat serum (Sigma-Aldrich), in 0.1% (v/v) Triton X-100 (Thermo Scientific) in PBS and cells were further incubated with primary antibodies overnight at 4°C, followed by incubation with Alexa– conjugated secondary antibodies. Before mounting (fluorescence mounting medium, S3023, Agilent Dako), neurons were counterstained with 300 nM DAPI in PBS (Sigma-Aldrich). For blocking peptide experiments, ten times excess of blocking peptide was incubated with antibody overnight at 4°C.

### Negative-stain transmission electron microscopy

Transmission electron microscopy projection images were collected using a JEM1400-Plus microscope (JEOL) operated at 100 kV and equipped with a Gatan OneView camera (4k 9 4k). High magnification (20,000x) and panoramic (5,000x) micrographs were obtained for synaptic vesicle and synapse density analyses, respectively. Only synapses with defined postsynaptic densities and containing between 10-200 synaptic vesicles, were included in the ultrastructural analysis. Synaptic vesicles and synapses were manually counted in blinded coded images to avoid bias. Synaptic vesicle diameter and presynaptic area were measured using ImageJ (*57*). For three-dimensional reconstructions, serial micrographs were aligned using Reconstruct software (https://synapseweb.clm.utexas.edu/software-0). Spatial frequency density plots for each synapse were generated by measuring vesicle coordinate positions and active zone structures based on electron micrographs oriented with the active zone at the bottom. For all synapses, vesicle positions were plotted on a grid aligned with respect to the center of the active zone and converted to a color-coded spatial density distribution. Vesicle positions assumed lateral symmetry around the midline (asymmetrical features are not informative because synapses are collected in all orientations in the slice). Euclidean distances from each vesicle to its nearest point on the active zone were calculated to generate cumulative distribution plots (*40*).

### Patch-clamp recording of cultured hippocampal neurons

Neurons were cultured on 13 mm coverslips as previously described (*56*). Immediately prior to recording, coverslips were transferred to a submerged-style recording chamber where they were continuously perfused (1-3 ml/min) with Carbogen (95% O_2_, 5% CO_2_), artificial cerebrospinal fluid (aCSF, 124 mM NaCl, 3 mM KCl, 24 mM NaHCO_3,_ 2 mM CaCl_2,_ 1.25 mM NaH_2_PO_4_, 1 mM MgSO_4_, 10 mM D-glucose) maintained at 34°C with a temperature control system (Scientifica, UK). Neurons were visualized with an infrared, differential interference contrast microscope (Scientifica, UK). Borosilicate glass micropipettes were pulled with a horizontal puller (Sutter, USA) to an access resistance of 5-9 MΩ. Single micropipettes were filled with intracellular solution (120 mM K-gluconate, 10 mM Na_2_-phosphocreatine, 0.3 mM Na_2_-GTP, 10 mM HEPES, 4 mM KCl, 4 mM Mg-ATP, pH 7.2, 280-290 mOsm). Some recordings were conducted in the presence of 30 µM Alexa Fluor^TM^ 488 dye in the intracellular solution. Following the clamping of a stable whole-cell configuration, the junction potential V_j_ = 15 mV arose due to the difference in composition between intracellular and extracellular solutions; this was corrected for arithmetically during analysis. All signals were amplified with a Multiclamp 700B amplifier and digitized using a Digidata 1550B. For whole cell recordings, a hyperpolarizing current of500 ms, −100 pA was injected, and the consequent plasma membrane voltage (V_m_) deflection was measured. To avoid the possibility of bias arising from cell-to-cell variability of the resting membrane potential, all recordings were conducted in the presence of a constant current, holding the pre-stimulus at V_m_ = −80 mV. For voltage-clamp recording of sEPSCs, a gap-free protocol at a holding potential V_h_ = −70 mV was used. Data analysis for quantifying the frequency and the waveform properties of sEPSCs used Clampfit (Molecular Devices, USA). The template search analysis tool embedded in Clampfit was used to identify single sEPSCs for each cell, with a manual check to identify any false positives and negatives. The average frequency was calculated as the number of events detected during 30 s, expressed in Hz. Averaged sEPSCs from all detected events were used to measure sEPSC waveform properties in each cell. Analysis tools in Clampfit were used to quantify the sEPSC rate of rise, peak, and halfwidth. Current-Voltage (I-V) curves for voltage-gated, non-inactivating, outward currents were carried out as follows. Series resistance was compensated for (10%-95% correction) and the capacitance of the pipette neutralized. Outward, non-inactivating, voltage gated currents were evoked by applying n=12, 30 ms, 10 mV voltage steps, starting from an initial V_h_ of −90 mV. Each recorded current was leak subtracted and normalized to membrane capacitance to measure the specific current. The specific conductance (G) was calculated as the ratio between the specific current and the electrochemical force for potassium (E_K_ = 100 mV). Cell-to-cell Boltzmann sigmoidal fit was used to estimate the maximal specific conductance (Gmax) and the half-activation voltage (V_½_) for each neuron. Current clamp recordings used a set pre-stimulus V_m_ of −80 mV, with a constant current injection (apart from recording the resting membrane potential). Passive electrical properties were measured by injecting a −100 pA, 500 ms square current step. The voltage deflection caused by this current injection followed a single exponential decay function. The steady state voltage deflection (A) and the voltage extrapolated *ad infinitum* (B) for a single exponential function fit between 10% and 90% of the minimum point (C) of the voltage deflection, were used to calculate cell input resistance (R_in_) and sag. Sag_sub_ was calculated as (C-A)/C and sag_fit_ was calculated as (B-A)/B, expressed as a percentage. Sag was measured as subtraction of the negative peak (sag_sub_) and of the extrapolated first exponential fit of the V_m_ decay (sag_fit_) upon injection of a hyperpolarizing step (*24*).

### Statistical analysis

Statistical analyses were performed using GraphPad Prism. Student’s t-test, one-way or two-way analysis of variance was used to determine differences between groups. Each experiment was performed at least in triplicate, as indicated.

## Acknowledgements

We thank the late Professor Peter Davies (Feinstein Institute for Medical Research, USA) for generously providing the PHF1 antibody used in this study. We thank the Wohl Cellular Imaging Centre at King’s College London for assistance with light microscopy. The authors acknowledge the Electron Microscopy Imaging Centre of the University of Sussex, funded by the School of Life Sciences, the Wellcome Trust (095605/Z/11/A and 208348/Z/17/Z) and the RM Phillips fund, for their support and assistance in this work. We thank the London Neurodegenerative Diseases Brain Bank at King’s College London for providing human post-mortem tissue.

## Author contributions

Conceptualization: DG, FT, DPH, KS.

Experimentation and data analysis: DG, FT, LB, SLR, SU, CT, SJP, HS, KS.

Manuscript writing and editing: DG, DPH, FT, KS, WN.

Supervision and funding: DPH, KS, LCS, WN.

## Data and materials availability

All data needed to evaluate the conclusions in the paper are present in the paper and/or the Supplementary Materials. Detailed numerical data are available on the data repository FigShare (DOI: https://figshare.com/s/295315cb2a4fde719963).

## Material availability

Materials are available upon reasonable request

## References

1. T. M. Wishart, S. H. Parson, T. H. Gillingwater, Synaptic vulnerability in neurodegenerative disease. J Neuropathol Exp Neurol 65, 733–739 (2006).

2. A. Kneynsberg, B. Combs, K. Christensen, G. Morfini, N. M. Kanaan, Axonal Degeneration in Tauopathies: Disease Relevance and Underlying Mechanisms. Front Neurosci 11, 572 (2017).

3. A. Lleo, R. Nunez-Llaves, D. Alcolea, C. Chiva, D. Balateu-Panos, M. Colom-Cadena, G. Gomez-Giro, L. Munoz, M. Querol-Vilaseca, J. Pegueroles, L. Rami, A. Llado, J. L. Molinuevo, M. Tainta, J. Clarimon, T. Spires-Jones, R. Blesa, J. Fortea, P. Martinez-Lage, R. Sanchez-Valle, E. Sabido, A. Bayes, O. Belbin, Changes in Synaptic Proteins Precede Neurodegeneration Markers in Preclinical Alzheimer’s Disease Cerebrospinal Fluid. Mol Cell Proteomics 18, 546–560 (2019).

4. C. M. Henstridge, B. T. Hyman, T. L. Spires-Jones, Beyond the neuron-cellular interactions early in Alzheimer disease pathogenesis. Nat Rev Neurosci 20, 94–108 (2019).

5. M. M. Shah, Neuronal HCN channel function and plasticity. Curr Opin Physiol 2, 92–97 (2018).

6. M. M. Shah, Cortical HCN channels: function, trafficking and plasticity. J Physiol 592, 2711–2719 (2014).

7. C. He, F. Chen, B. Li, Z. Hu, Neurophysiology of HCN channels: from cellular functions to multiple regulations. Prog Neurobiol 112, 1–23 (2014).

8. Z. Huang, G. Li, C. Aguado, R. Lujan, M. M. Shah, HCN1 channels reduce the rate of exocytosis from a subset of cortical synaptic terminals. Sci Rep 7, 40257 (2017).

9. C. D. Paspalas, M. Wang, A. F. Arnsten, Constellation of HCN channels and cAMP regulating proteins in dendritic spines of the primate prefrontal cortex: potential substrate for working memory deficits in schizophrenia. Cereb Cortex 23, 1643–1654 (2013).

10. J. C. Magee, Dendritic Ih normalizes temporal summation in hippocampal CA1 neurons. Nat Neurosci 2, 848 (1999).

11. D. Tsay, J. T. Dudman, S. A. Siegelbaum, HCN1 channels constrain synaptically evoked Ca2+ spikes in distal dendrites of CA1 pyramidal neurons. Neuron 56, 1076–1089 (2007).

12. Z. Huang, R. Lujan, I. Kadurin, V. N. Uebele, J. J. Renger, A. C. Dolphin, M. M. Shah, Presynaptic HCN1 channels regulate Cav3.2 activity and neurotransmission at select cortical synapses. Nat Neurosci 14, 478–486 (2011).

13. X. Chang, J. Wang, H. Jiang, L. Shi, J. Xie, Hyperpolarization-Activated Cyclic Nucleotide-Gated Channels: An Emerging Role in Neurodegenerative Diseases. Front Mol Neurosci 12, 141 (2019).

14. T. F. Musial, E. Molina-Campos, L. A. Bean, N. Ybarra, R. Borenstein, M. L. Russo, E. W. Buss, D. Justus, K. M. Neuman, G. D. Ayala, S. A. Mullen, Y. Voskobiynyk, C. T. Tulisiak, J. A. Fels, N. J. Corbett, G. Carballo, C. D. Kennedy, J. Popovic, J. Ramos-Franco, M. Fill, M. R. Pergande, J. A. Borgia, G. T. Corbett, K. Pahan, Y. Han, D. M. Chetkovich, R. J. Vassar, R. W. Byrne, M. Matthew Oh, T. R. Stoub, S. Remy, J. F. Disterhoft, D. A. Nicholson, Store depletion-induced h-channel plasticity rescues a channelopathy linked to Alzheimer’s disease. Neurobiol Learn Mem 154, 141–157 (2018).

15. J. L. Crimins, A. B. Rocher, J. I. Luebke, Electrophysiological changes precede morphological changes to frontal cortical pyramidal neurons in the rTg4510 mouse model of progressive tauopathy. Acta Neuropathol 124, 777–795 (2012).

16. C. A. Booth, T. Ridler, T. K. Murray, M. A. Ward, E. de Groot, M. Goodfellow, K. G. Phillips, A. D. Randall, J. T. Brown, Electrical and Network Neuronal Properties Are Preferentially Disrupted in Dorsal, But Not Ventral, Medial Entorhinal Cortex in a Mouse Model of Tauopathy. J Neurosci 36, 312–324 (2016).

17. M. K. Bondulich, T. Guo, C. Meehan, J. Manion, T. Rodriguez Martin, J. C. Mitchell, T. Hortobagyi, N. Yankova, V. Stygelbout, J. P. Brion, W. Noble, D. P. Hanger, Tauopathy induced by low level expression of a human brain-derived tau fragment in mice is rescued by phenylbutyrate. Brain : a journal of neurology 139, 2290–2306 (2016).

18. O. Shupliakov, V. Haucke, A. Pechstein, How synapsin I may cluster synaptic vesicles. Semin Cell Dev Biol 22, 393–399 (2011).

19. R. Yuste, T. Bonhoeffer, Morphological changes in dendritic spines associated with long-term synaptic plasticity. Annu Rev Neurosci 24, 1071–1089 (2001).

20. K. P. Berry, E. Nedivi, Spine Dynamics: Are They All the Same? Neuron 96, 43–55 (2017).

21. J. Herms, M. M. Dorostkar, Dendritic Spine Pathology in Neurodegenerative Diseases. Annu Rev Pathol 11, 221–250 (2016).

22. J. Roos, R. B. Kelly, The endocytic machinery in nerve terminals surrounds sites of exocytosis. Curr Biol 9, 1411–1414 (1999).

23. V. I. Slepnev, P. De Camilli, Accessory factors in clathrin-dependent synaptic vesicle endocytosis. Nat Rev Neurosci 1, 161–172 (2000).

24. F. Tamagnini, S. Scullion, J. T. Brown, A. D. Randall, Intrinsic excitability changes induced by acute treatment of hippocampal CA1 pyramidal neurons with exogenous amyloid beta peptide. Hippocampus 25, 786–797 (2015).

25. H. Richter, U. Heinemann, C. Eder, Hyperpolarization-activated cation currents in stellate and pyramidal neurons of rat entorhinal cortex. Neurosci Lett 281, 33–36 (2000).

26. S. J. Barnes, C. E. Cheetham, Y. Liu, S. H. Bennett, G. Albieri, A. A. Jorstad, G. W. Knott, G. T. Finnerty, Delayed and Temporally Imprecise Neurotransmission in Reorganizing Cortical Microcircuits. J Neurosci 35, 9024–9037 (2015).

27. N. Chuhma, H. Ohmori, Postnatal development of phase-locked high-fidelity synaptic transmission in the medial nucleus of the trapezoid body of the rat. J Neurosci 18, 512–520 (1998).

28. N. L. Chanaday, M. A. Cousin, I. Milosevic, S. Watanabe, J. R. Morgan, The Synaptic Vesicle Cycle Revisited: New Insights into the Modes and Mechanisms. J Neurosci 39, 8209–8216 (2019).

29. T. Spires-Jones, S. Knafo, Spines, plasticity, and cognition in Alzheimer’s model mice. Neural Plast 2012, 319836 (2012).

30. O. Franz, B. Liss, A. Neu, J. Roeper, Single-cell mRNA expression of HCN1 correlates with a fast gating phenotype of hyperpolarization-activated cyclic nucleotide-gated ion channels (Ih) in central neurons. Eur J Neurosci 12, 2685–2693 (2000).

31. L. M. Giocomo, S. A. Hussaini, F. Zheng, E. R. Kandel, M. B. Moser, E. I. Moser, Grid cells use HCN1 channels for spatial scaling. Cell 147, 1159–1170 (2011).

32. G. M. F. Nolan, G. Malleret, J. T. Dudman, D. L. Buhl, B. Santoro, E. Gibbs, S. Vronskaya, G. Buzsaki, S. A. Siegelbaum, E. R. Kandel, A. Morozov, A behavioral role for dendritic integration: HCN1 channels constrain spatial memory and plasticity at inputs to distal dendrites of CA1 pyramidal neurons. Cell 119, 719–732 (2004).

33. J. S. Jackson, J. Witton, J. D. Johnson, Z. Ahmed, M. Ward, A. D. Randall, M. L. Hutton, J. T. Isaac, M. J. O’Neill, M. C. Ashby, Altered Synapse Stability in the Early Stages of Tauopathy. Cell Rep 18, 3063–3068 (2017).

34. C. A. Booth, J. Witton, J. Nowacki, K. Tsaneva-Atanasova, M. W. Jones, A. D. Randall, J. T. Brown, Altered Intrinsic Pyramidal Neuron Properties and Pathway-Specific Synaptic Dysfunction Underlie Aberrant Hippocampal Network Function in a Mouse Model of Tauopathy. J Neurosci 36, 350–363 (2016).

35. M. Ludwig, G. Leng, Dendritic peptide release and peptide-dependent behaviours. Nat Rev Neurosci 7, 126–136 (2006).

36. J. Marksteiner, W. A. Kaufmann, P. Gurka, C. Humpel, Synaptic proteins in Alzheimer’s disease. J Mol Neurosci 18, 53–63 (2002).

37. K. R. Christensen, T. G. Beach, G. E. Serrano, N. M. Kanaan, Pathogenic tau modifications occur in axons before the somatodendritic compartment in mossy fiber and Schaffer collateral pathways. Acta Neuropathol Commun 7, 29 (2019).

38. D. Milovanovic, Y. Wu, X. Bian, P. De Camilli, A liquid phase of synapsin and lipid vesicles. Science 361, 604–607 (2018).

39. R. Sansevrino, C. Hoffmann, D. Milovanovic, Condensate biology of synaptic vesicle clusters. Trends Neurosci, (2023).

40. A. Orenbuch, L. Shalev, V. Marra, I. Sinai, Y. Lavy, J. Kahn, J. J. Burden, K. Staras, D. Gitler, Synapsin selectively controls the mobility of resting pool vesicles at hippocampal terminals. J Neurosci 32, 3969–3980 (2012).

41. V. Marra, J. J. Burden, J. R. Thorpe, I. T. Smith, S. L. Smith, M. Hausser, T. Branco, K. Staras, A preferentially segregated recycling vesicle pool of limited size supports neurotransmission in native central synapses. Neuron 76, 579–589 (2012).

42. S. Rey, V. Marra, C. Smith, K. Staras, Nanoscale Remodeling of Functional Synaptic Vesicle Pools in Hebbian Plasticity. Cell Rep 30, 2006–2017 e2003 (2020).

43. S. A. Rey, C. A. Smith, M. W. Fowler, F. Crawford, J. J. Burden, K. Staras, Ultrastructural and functional fate of recycled vesicles in hippocampal synapses. Nat Commun 6, 8043 (2015).

44. C. Wahl-Schott, M. Biel, HCN channels: structure, cellular regulation and physiological function. Cell Mol Life Sci 66, 470–494 (2009).

45. Y. Saito, T. Inoue, G. Zhu, N. Kimura, M. Okada, M. Nishimura, N. Kimura, S. Murayama, S. Kaneko, R. Shigemoto, K. Imoto, T. Suzuki, Hyperpolarization-activated cyclic nucleotide gated channels: a potential molecular link between epileptic seizures and Abeta generation in Alzheimer’s disease. Mol Neurodegener 7, 50 (2012).

46. T. E. Chater, Y. Goda, The role of AMPA receptors in postsynaptic mechanisms of synaptic plasticity. Front Cell Neurosci 8, 401 (2014).

47. F. Tamagnini, D. A. Walsh, J. T. Brown, M. K. Bondulich, D. P. Hanger, A. D. Randall, Hippocampal neurophysiology is modified by a disease-associated C-terminal fragment of tau protein. Neurobiology of aging 60, 44–56 (2017).

48. H. S. Wang, Z. Pan, W. Shi, B. S. Brown, R. S. Wymore, I. S. Cohen, J. E. Dixon, D. McKinnon, KCNQ2 and KCNQ3 potassium channel subunits: molecular correlates of the M-channel. Science 282, 1890–1893 (1998).

49. S. L. Schmidt, C. R. Dorsett, A. K. Iyengar, F. Frohlich, Interaction of Intrinsic and Synaptic Currents Mediate Network Resonance Driven by Layer V Pyramidal Cells. Cereb Cortex 27, 4396–4410 (2017).

50. Y. Buskila, O. Kekesi, A. Bellot-Saez, W. Seah, T. Berg, M. Trpceski, J. J. Yerbury, L. Ooi, Dynamic interplay between H-current and M-current controls motoneuron hyperexcitability in amyotrophic lateral sclerosis. Cell Death Dis 10, 310 (2019).

51. R. Hu, K. A. Ferguson, C. B. Whiteus, D. H. Meijer, R. C. Araneda, Hyperpolarization-Activated Currents and Subthreshold Resonance in Granule Cells of the Olfactory Bulb. eNeuro 3, (2016).

52. A. Boehlen, C. Henneberger, U. Heinemann, I. Erchova, Contribution of near-threshold currents to intrinsic oscillatory activity in rat medial entorhinal cortex layer II stellate cells. J Neurophysiol 109, 445–463 (2013).

53. H. Hu, K. Vervaeke, J. F. Storm, Two forms of electrical resonance at theta frequencies, generated by M-current, h-current and persistent Na+ current in rat hippocampal pyramidal cells. J Physiol 545, 783–805 (2002).

54. G. G. Turrigiano, S. B. Nelson, Homeostatic plasticity in the developing nervous system. Nat Rev Neurosci 5, 97–107 (2004).

55. G. F. Kusick, M. Chin, S. Raychaudhuri, K. Lippmann, K. P. Adula, E. J. Hujber, T. Vu, M. W. Davis, E. M. Jorgensen, S. Watanabe, Synaptic vesicles transiently dock to refill release sites. Nat Neurosci, (2020).

56. C. Giacomini, C. Y. Koo, N. Yankova, I. A. Tavares, S. Wray, W. Noble, D. P. Hanger, J. D. H. Morris, A new TAO kinase inhibitor reduces tau phosphorylation at sites associated with neurodegeneration in human tauopathies. Acta Neuropathol Commun 6, 37 (2018).

57. C. A. Schneider, W. S. Rasband, K. W. Eliceiri, NIH Image to ImageJ: 25 years of image analysis. Nat Methods 9, 671–675 (2012).

