## Supplementary Figures for "Tau-mediated synaptic dysfunction is coupled with HCN channelopathy"

Despoina Goniotaki *et al.*

**This section includes:**

Supplementary Text
Figs. S1 to S5

### Supplementary Text

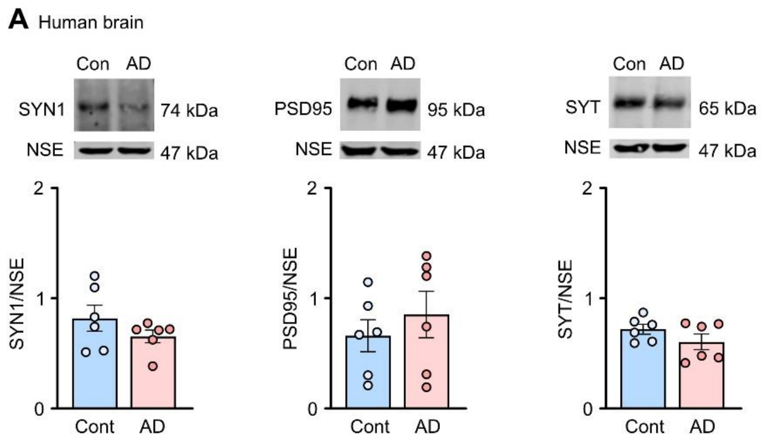


### Fig. S1. Analysis of synaptic markers in human postmortem AD tissue

(**A**) Western blots of brain homogenates from Alzheimer's disease (AD) and control postmortem hippocampus probed with antibodies to synapsin 1 (SYN1), postsynaptic density 95 (PSD95), synaptotagmin (SYT) and neuron specific enolase (NSE). Quantifications are shown in the graphs as mean ± SEM; n= 6 brains per group. Student's t-test.

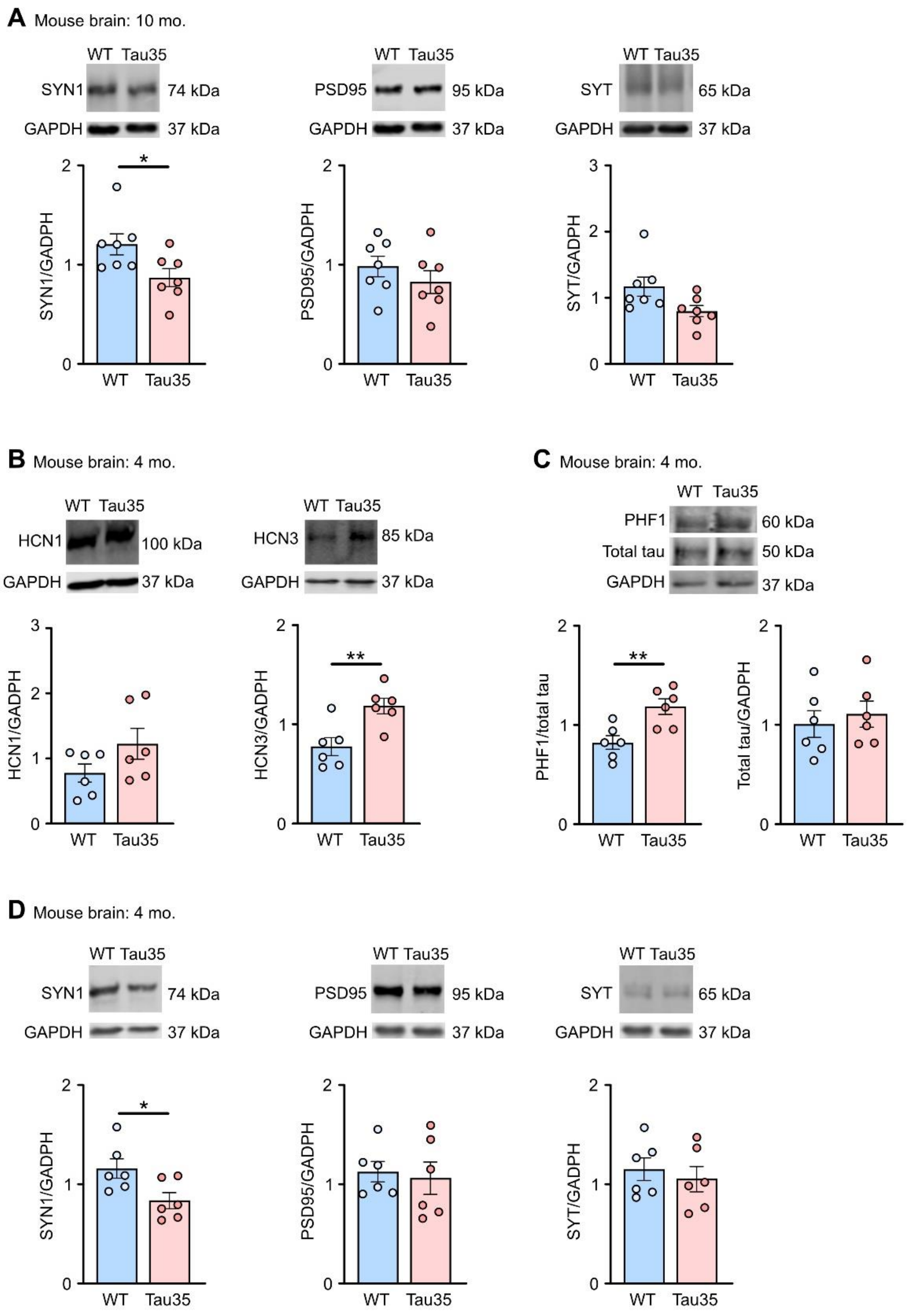
A Mouse brain: 10 mo.
WT Tau35
SYN1 74 kDa
GAPDH 37 kDa
PSD95 95 kDa
SYT 65 kDa
SYN1/GADPH
PSD95/GADPH
SYT/GADPH
WT Tau35
B Mouse brain: 4 mo.
HCN1 100 kDa
HCN3 85 kDa
HCN1/GADPH
HCN3/GADPH
C Mouse brain: 4 mo.
PHF1 60 kDa
Total tau 50 kDa
PHF1/total tau
Total tau/GADPH
D Mouse brain: 4 mo.
*
**

### Fig. S2. Analysis of synaptic markers in Tau35 and wild-type (WT) brain homogenates

(**A**) Western blots of brain homogenates from wild-type (WT) and Tau35 mice aged 10 months, probed with antibodies to synapsin 1 (SYN1), synaptotagmin (SYT), postsynaptic density 95 (PSD95) and glyceraldehyde 3-phosphate dehydrogenase (GAPDH). Quantification of the blots is shown in the graphs as mean ± SEM, n= 6-7 brains per group. Student's t-test, $*P < 0.05$.

(**B to D**) Western blots of brain homogenates from wild-type (WT) and Tau35 mice aged 4 months (pre-symptomatic) probed with antibodies to hyperpolarisation-activated cyclic nucleotide-gated (HCN) channels (HCN1 and HCN3), phosphorylated tau (PHF1), total tau, synapsin 1 (SYN1), synaptotagmin (SYT), postsynaptic density 95 (PSD95) and glyceraldehyde 3-phosphate dehydrogenase (GAPDH). Quantification of the blots is shown in the graphs as mean ± SEM; n= 6-7 brains per group. Student's t-test, $*P < 0.05$, $**P < 0.01$.

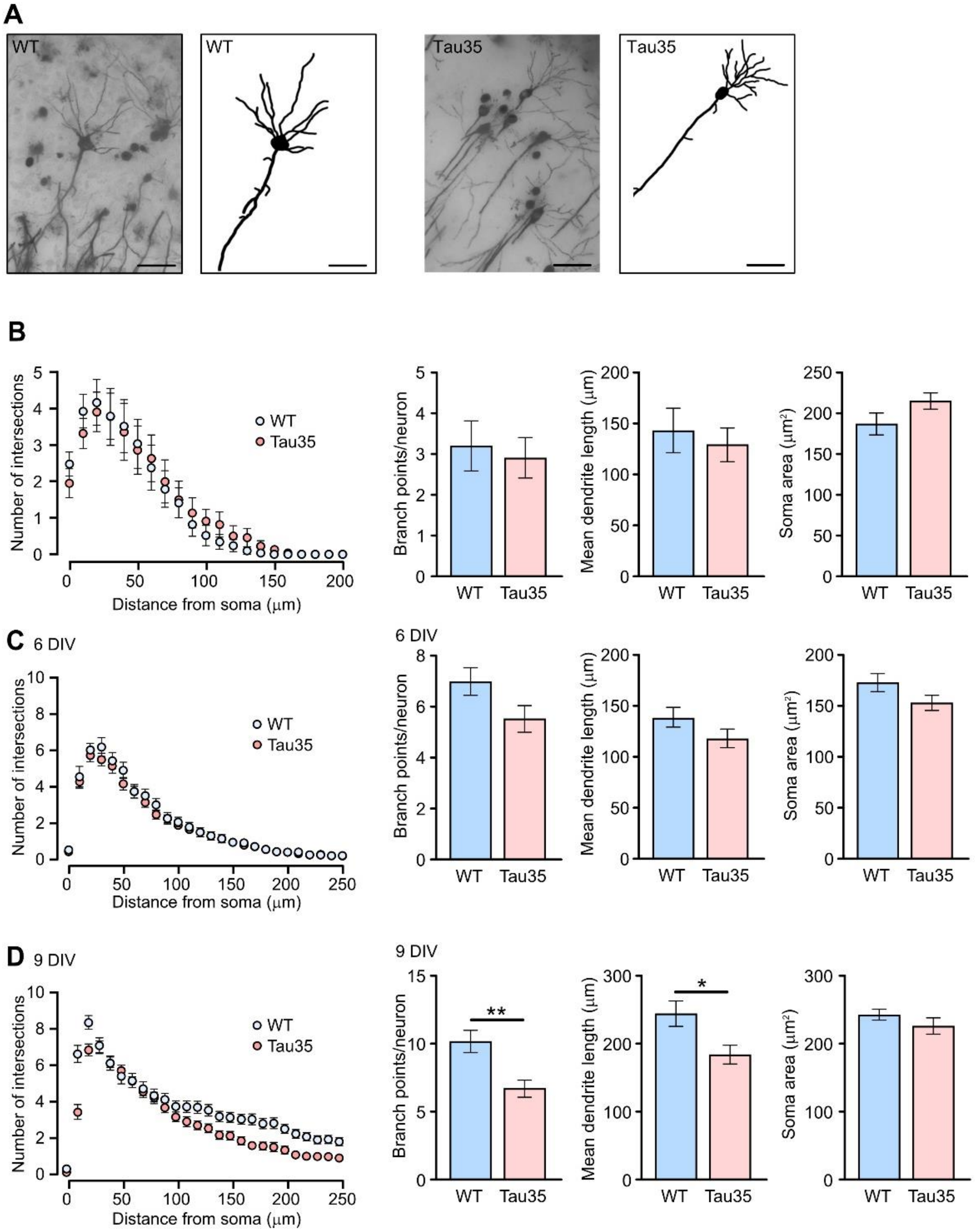


### Fig. S3. Dendritic branching in young Tau35 mice and Tau35 hippocampal neurons

(**A**) Images of Golgi-Cox stained wild-type (WT) and Tau35 CA1 hippocampal neurons in the brains of 4-month-old mice. Scale bars: 50μm. (**B**) Total Sholl analysis shows no differences in basal dendrite complexity in neurons of WT and Tau35 mice aged 4 months. Graph shows quantification of the mean ± SEM, n=40 neurons from 4-5 mice of each genotype. Two-way ANOVA, $P > 0.05$. Graphs show the number of primary branch points, dendrite length and soma area in neurons of WT and Tau35 mice aged 4 months, n=40 neurons from 4-5 mice of each genotype. Student's t-test, $P > 0.05$. (**C**) Sholl analysis of dendritic branching in wild-type (WT) and Tau35 primary hippocampal neurons at 6 DIV. Graph shows mean ± SEM. Two-way ANOVA ($P < 0.01$), n=50 neurons of each genotype from 5 independent experiments. Graphs below show mean ± SEM number of dendritic branch points, mean dendrite length and soma area, n=39-52 neurons from 4-5 independent experiments. Student's t-test, $P > 0.05$. (**D**) Sholl analysis of dendritic branching in WT and Tau35 hippocampal neurons at 9 DIV. Graph shows mean ± SEM. Two-way ANOVA ($P < 0.01$), n=55 neurons of each genotype from 5 independent experiments. Graphs below show mean ± SEM of the number of dendritic branch points, mean dendrite length and soma area, n=55 neurons of each genotype from 5 independent experiments. Student's t-test, *$P < 0.05$, **$P < 0.01$.

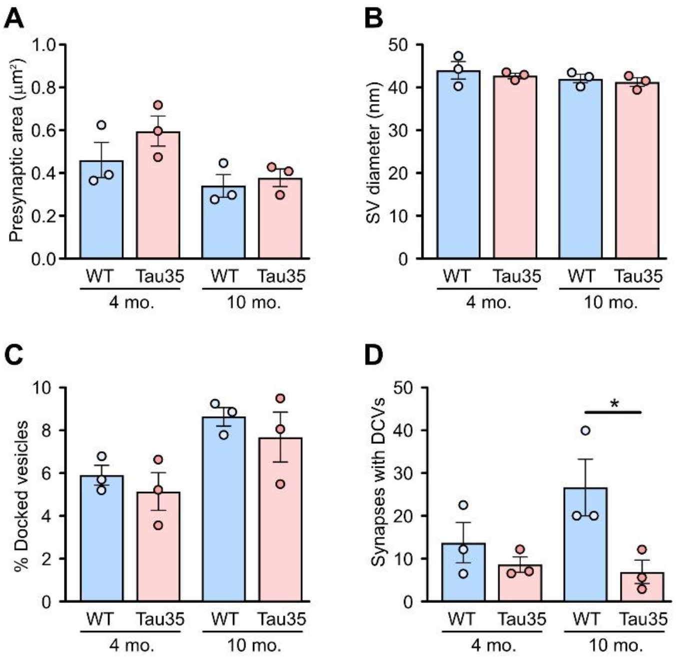


**Fig. S4. Ultrastructural analysis of wild-type and Tau35 hippocampal synapses**

(**A to C**) Quantification of SV diameter and presynaptic area of synapses in the CA1 region of the hippocampus in WT and Tau35 mice aged 4 and 10 months. **c** Graphs show the percentage of docked vesicles/synapse in WT and Tau35 mice aged 4 and 10 months. (**D**) Graphs show the percentage of synapses harbouring DCVs in WT and Tau35 mice aged 4 and 10 months. For (**A to D**): Graphs show mean ± SEM, n=100-150 synapses from 3 mice of each genotype. Two-way ANOVA, *$P < 0.05$.

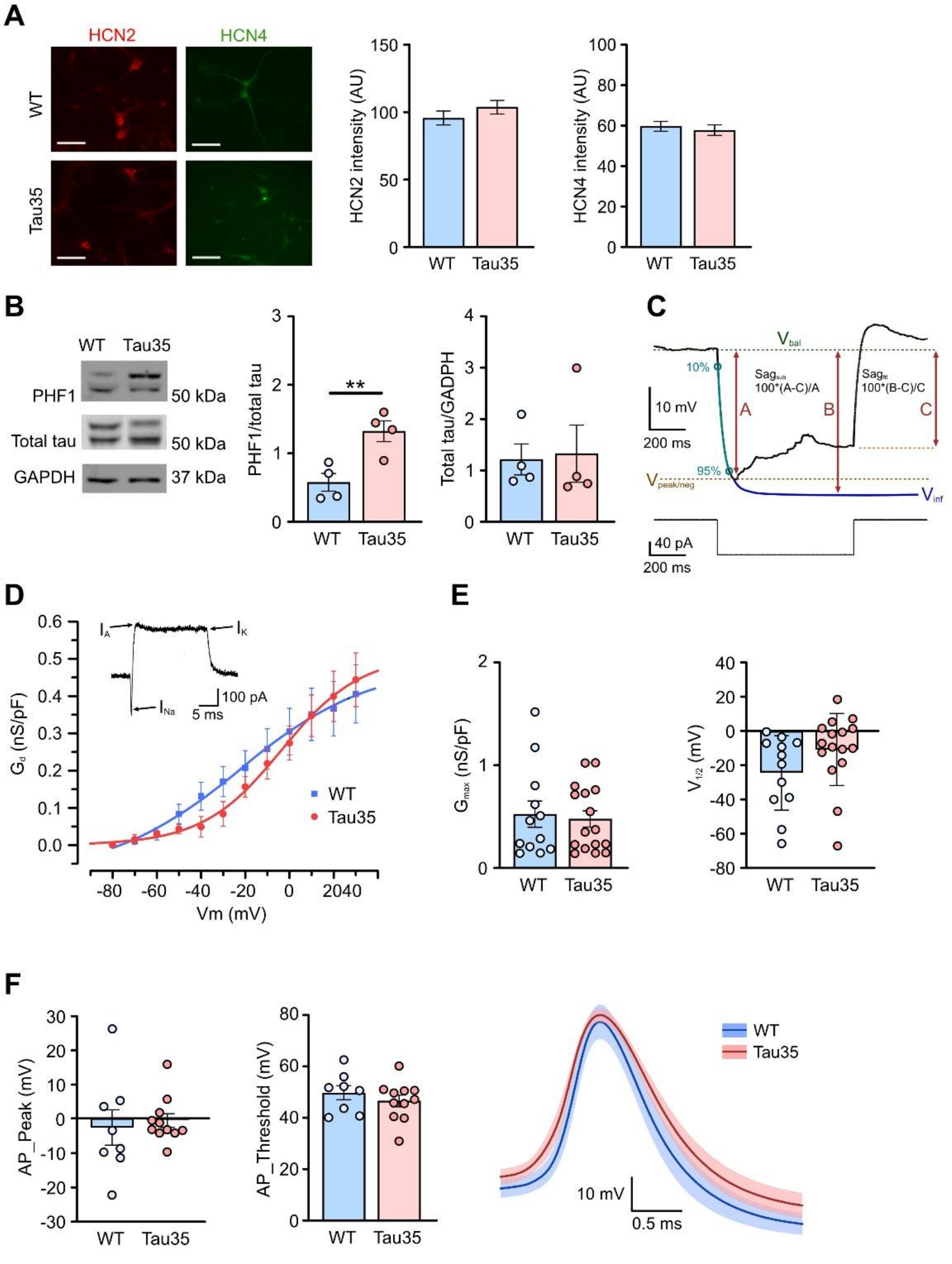

### Fig S5. $I_k$ conductance and action potential characteristics in Tau35 hippocampal neurons

(**A**) Immunofluorescence labelling of WT and Tau35 mouse hippocampal neurons at 14 DIV with HC2 and HCN4 antibodies. Scale bars: 50μm. Graphs show quantification of mean fluorescence intensity (± SEM) of HCN2 and HCN4. n=150 (WT) and n=120 (Tau35) neurons, from 3 independent experiments. Student's t-test. (**B**) Western blots of lysates of primary hippocampal neurons (14 DIV) from wild-type (WT) and Tau35 mice, probed with antibodies to phosphorylated tau (PHF1) and total tau and glyceraldehyde 3-phosphate dehydrogenase (GAPDH). Quantification of the blots is shown in the graphs as mean ± SEM; n=4 independent experiments. Student's t-test, $**P < 0.01$. (**C**) Schematic representation of the deflection of the plasma membrane voltage in response to a hyperpolarizing current step (500 ms, -100 pA) to explore $I_h$-dependent sag properties. The steady state voltage deflection (A) and the voltage extrapolated *ad infinitum* (B) for a single exponential function fit between 10% and 90% of the minimum point (C) of the voltage deflection, were used to calculate cell input resistance ($R_{in}$) and sag. $Sag_{sub}$ was calculated as (A-C)/A and $sag_{fit}$ was calculated as (B-C)/B and expressed as a percentage. (**D**) $I_k$ conductance in wild-type (WT) and Tau35 mouse hippocampal neurons (11-16 DIV). Average ± SEM values are shown in relation to different voltage steps. The example trace in the top insert, evoked by a depolarizing voltage step (30 ms) reveals the different voltage gated components evoked by depolarization. (**E**) Graphs showing no effect of genotype on either the maximal conductance (Gmax) or the half-activation potential ($V_{1/2}$). Graphs show mean ± SEM, Student's t-test, n=12-16 neurons from 3 independent experiments. (**F**) AP-peak (amplitude) (mV) and AP-threshold (mV) of the plasma membrane and comparative action potential (AP) waveforms of WT and Tau35 neurons at 11-16 DIV. Graphs show mean ± SEM, n=8-11 neurons from 3 independent experiments. Student's t-test, $P > 0.05$.
